# GABAergic neuronal IL-4R mediates T cell effect on memory

**DOI:** 10.1101/2021.03.16.435727

**Authors:** Zhongxiao Fu, Taitea Dykstra, Morgan Wall, Huiping Li, Andrea Francesca Salvador, Bende Zou, Ni Yan, Patrick H. Andrews, Dylan H. Goldman, Zachary Papadopoulos, Igor Smirnov, Xinmin S. Xie, Jasmin Herz, Jonathan Kipnis

## Abstract

Mechanisms governing how immune cells and their derived molecules impact homeostatic brain function are still poorly understood. Here, we elucidate neuronal mechanisms underlying effects of T cells on synaptic function and episodic memory. Depletion of CD4 T cells led to memory deficits and impaired long term potentiation. Severe combined immune-deficient (SCID) mice exhibited amnesia, which was reversible by repopulation with T cells from wild-type but not from IL-4-knockout mice. This rescue was mediated via IL-4 receptors (IL-4R) expressed on neurons. Exploration of snRNAseq of neurons participating in memory processing provided insights into synaptic organization and plasticity-associated pathways and genes regulated by immune cells and molecules. IL-4Rα knockout in inhibitory neurons impaired contextual fear memory, suggesting participation of an IL-4-associated switch in regulating synaptic function and promoting contextual fear memory. These findings provide insights into neuroimmune interactions at the transcriptional and functional levels in neurons.

## Introduction

The nervous system integrates ongoing environment-derived information with previously stored experiences, thereby guiding behaviors and allowing environmental adaptation. Concomitantly the immune system, based on experience-derived memory, protects the host against microorganisms, allowing its better adaptation and survival. Despite similarities in principles guiding both systems, their communication in support of host adaptation is still elusive.

Episodic memory is a recollection of past experiences conveying distinct temporal and spatial information (Tulving, 2002). In the processing of such “mental time travel”, both in animal models and in humans, the hippocampus has a key role (Bird and Burgess, 2008; Burgess et al., 2002; Eichenbaum et al., 1999; Frisk and Milner, 1990; Hitti and Siegelbaum, 2014). The mechanisms of episodic memory have been dissected in numerous studies at multiple levels, including actively involved brain regions (Chen et al., 1996; Frisk and Milner, 1990), neuronal circuits (Herry et al., 2008; Roy et al., 2017), associated synaptic strength changes (Mahan and Ressler, 2012; McKernan and Shinnick-Gallagher, 1997), and molecular mechanisms (Johansen et al., 2011).

The immune system has been linked to cognitive function in a series of studies (Baruch et al., 2014; Castellano et al., 2017; Derecki et al., 2010a; Kipnis et al., 2004; Villeda et al., 2014), mostly employing complex prolonged behavioral tasks in devices such as the Morris water maze (MWM), albeit without adequate understanding of the mechanisms invoked. Immune-deficient mice exhibited impaired spatial memory performance in the MWM, and reconstitution of splenocytes reversed the cognitive deficits (Derecki et al., 2010b). The underlying mechanism(s) were not addressed, although similar spatial memory deficits were observed in mice deficient in interleukin (IL)-4 or IL-13, and an elusive mechanism mediated through astrocytes was suggested to account for it (Brombacher et al., 2017; Derecki et al., 2010b). However, whether and how T cells and their derived cytokines affect episodic memory processing is not yet known.

A recently emerging new line of evidence suggests that cytokines can act as neuromodulators, thereby possibly altering neuronal responses and modifying neuronal function by signaling via their neuronally expressed cognate receptors (Chen et al., 2017; Filiano et al., 2016a) (Choi et al., 2016; de Lima et al., 2020). This intriguing novel indication of cytokine-induced potential communication between the immune and the nervous systems prompted us to revisit the roles of T cells and their derived IL-4 in episodic memory. Our aim was to acquire a better mechanistic insight into whether and how episodic memory is promoted by IL-4.

Here, we show that contextual fear conditioning (CFC) memory is impaired after depletion of CD4 T cells in wild-type (WT) mice. Consistently with the behavioral findings, CD4 T-cell depletion also impaired long-term synaptic potentiation in the hippocampal dentate gyrus (DG). We further found that memory deficits in immune-deficient mice could be restored by wild type T cells but not by T cells deficient in IL-4, and that conditional knockout of IL-4Rα from neurons, but not from microglia, recapitulated memory deficits. Single-nuclei RNA sequencing (snRNAseq) of neurons responding to a memory task revealed that compared to WT, immune-deficient mice exhibited dysregulation of synapse-related gene expression, and that this can be largely reversed by repopulation with T cells. Our study thus shows that CD4 T cells and their derived IL-4 regulate episodic memory, providing a mechanistic view that T cells and IL-4 alter synaptic transmission, synaptic plasticity and gene expression in task-activated neurons.

## Results

### T cells regulate episodic memory through interleukin-4

To explore the potential neuro-immune interactions during episodic memory processing, we employed the CFC memory paradigm, a robust memory task with well-studied circuitry (Goshen et al., 2011; Liu et al., 2012b; Maren, 2001a) (**Figure 1A**). In this task, mice are introduced to a new environment (new context) and concomitantly exposed to an unconditioned stimulus of foot shock (Maren, 2001b; Phillips and LeDoux, 1992). They are then moved to a home cage for 1 day, and upon reintroduction to the “shock context” but without the stimulus, their immobility (freezing) in anticipation of a shock is quantified. The time that freezing lasts is indicative of contextual memory. We first determined whether CD4 T cells and IL-4 are required for CFC memory regulation. Acute depletion of CD4 T cells from WT mice (**Supplemental Figure S1A−C**) for 3 weeks resulted in their reduced freezing times during the memory test when compared to mice injected with control antibodies (**Figure 1B**). Long-term potentiation (LTP) via a theta-burst stimulation paradigm resulted in a significantly lower population spike amplitude (PSA) in CD4 T cell-depleted mice than in the control group (**Figure 1C, D**), supporting the notion that mice lacking T cells show impaired contextual fear memory and neuronal function.

**Figure 1:**
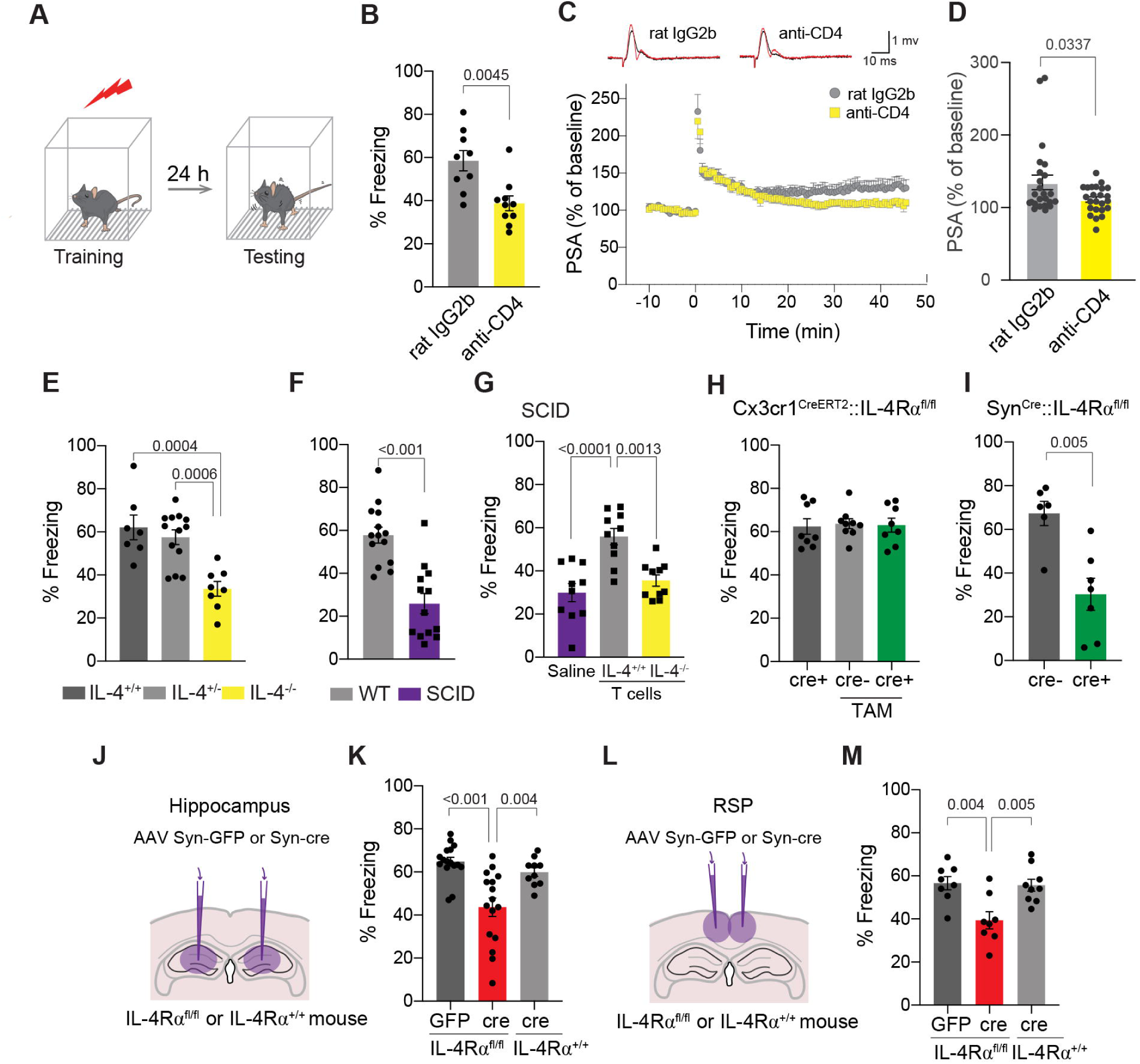
T cell-derived IL-4 promotes contextual fear memory through neuronal IL-4Rα. **A**, Schematic diagram of the behavioral experimental procedure. Mice were trained with contextual fear conditioning and subjected to the same cage (context) for testing 24 hours later. Percent freezing time during 3 minutes of testing. **B**, Percent freezing time of mice depleted of CD4 T cells. Mice were injected i.p. with either 200 ug rat IgG2b isotype control (clone LTF-2, n = 9 mice) or anti-CD4 antibody (clone GK1.5, n = 10 mice) followed by 100 ug weekly injections for three weeks. Two-tailed unpaired Mann-Whitney *U*-test. Representative experiment is shown out of 2 independent experiments performed. **C**, Long-term potentiation of population spike amplitude (PSA) in hippocampal dentate gyrus slices after CD4 T cell depletion. Upper panel: representative traces of WT mice treated with control rat IgG2b or anti-CD4 antibody for three weeks. Lower panel: Average traces of PSA obtained before and after LTP induction by theta-burst stimulation. N = 24 slices from 8 mice per group. **D,** Normalized PSA (% of baseline) for the last 5 minutes. Two-tailed unpaired Mann-Whitney *U*-test. **E,** Percent freezing time of IL-4^+/+^ (n=7), IL-4^-/+^ (n=13) and IL-4^-/-^ (n=8) mice. One-Way ANOVA with Tukey post hoc test. Two independent experiments combined are presented. **F**, Percent freezing time of WT and SCID mice 24 hours (n = 14 WT, n = 13 SCID) after training. Two-tailed unpaired Mann-Whitney *U*-test. Representative experiments are shown out of 3 independent experiments performed. **G**, Percent freezing time of SCID mice injected with saline, 2 x 10^6^ splenic IL4^+/+^ or IL4^-/-^ T cells. Mice underwent training and fear memory testing 4 weeks after repopulation (n = 10 mice per group). One-Way ANOVA with Tukey post hoc test. Representative experiment is shown out of 2 independent experiments performed. **H**, Percent freezing time of mice with microglia-specific knockout of IL-4Rα (n= 8-9 mice per group) and **I,** neuron-specific knockout of IL-4Rα (n= 6-7 mice per group) for 24-hour contextual fear memory testing. Two-tailed unpaired Mann-Whitney *U*-test. **J**, Schematic showing bilateral injection of AAV-Syn-GFP or AAV-Syn-cre virus into the hippocampus of IL-4Rα^+/+^ or IL-4Rα^fl/fl^ mice 4 weeks prior to behavioral testing. **K**, Percent freezing time for 24 hours testing of IL-4Rα^fl/fl^ mice injected with AAV-Syn-GFP (n = 16) or AAV-Syn-cre (n = 16) and IL-4Rα^+/+^ mice injected with AAV-Syn-cre (n = 10) into the hippocampus. One-Way ANOVA with Tukey post hoc test. **L,** Schematic showing bilateral injection of AAV-Syn-GFP or AAV-Syn-cre virus into retrosplenial cortex of IL-4Rα^+/+^ or IL-4Rα^fl/fl^ mice 4 weeks prior to behavioral testing. **M**, Percent freezing time for 24 hours testing of IL-4Rα^fl/fl^ mice injected with AAV-Syn-GFP (n = 8) or AAV-Syn-cre (n = 8) and IL-4Rα^+/+^ mice injected with AAV-Syn-cre (n = 9) into the retrosplenial cortex. One-Way ANOVA with Tukey post hoc test. Data are presented as means ± SEM.

Similar results were obtained in mice deficient in IL-4, a cytokine produced by Th2 lymphocytes (Seder et al, 1994). While control and IL-4 knockout mice showed similar freezing times before training (**Supplemental Figure S1D**), the freezing response of the IL-4 knockout mice during testing was impaired compared to that of their IL-4^−/+^ or IL-4^+/+^ littermates (**Figure 1E**). In addition, given that severe combined immune-deficient (SCID) mice lack adaptive immunity, we also examined the performance of the SCID mice in the memory task. WT and SCID mice showed similar freezing behavior when assessed before and 1 hr after training (**Supplemental Figure S1E, F**); however compared to their WT counterparts, significantly reduced freezing times were evident in both male and female SCID mice when assessed at 24 hr and at 4 weeks post-training (**Figure 1F**; **Supplemental Figure S1F, G**).

To find out whether memory deficit is a result of immunodeficiency, we adoptively transferred T cells from WT donors or from IL4^−/−^ donors into young adult SCID mice, in which CFC memory was tested 4 weeks after reconstitution. Compared to their non-repopulated counterparts (**Figure 1G**), the SCID mice showed significantly longer freezing behavior when repopulated with T cells from WT but not from IL-4^−/−^ mice, suggesting that T cells and their derived IL-4 are required for normal CFC memory. As expected, immunosufficient WT mice showed no changes in freezing behavior when repopulated with T cells from either WT or IL-4^−/−^ mice (**Supplemental Figure S1H**).

### IL-4 regulates CFC memory through neuronal IL-4R**α**

To better understand the IL-4-signaling processes underlying these changes, IL-4Rα was conditionally deleted from microglia. No memory deficits were observed (**Figure 1H**), however, conditional deletion of IL-4Rα from neurons resulted in impaired memory function (**Figure 1I**). Confirmation of IL-4Rα deletion from neurons (both GABAergic and glutamatergic) was assessed by RNA-fluorescence *in situ* hybridization (**Supplemental Figure S2C)**. To further explore the regional specificity of IL-4 function, we deleted IL-4Rα from hippocampal or retrosplenial cortex (RSP) neurons via stereotactic AAV-SynapsinI-cre delivery in IL-4Rα^fl/fl^ mice, and tested the mice for freezing behavior 24 hr after fear training. We found that deletion of IL-4Rα from hippocampal (**Figure 1J, K**) or RSP neurons (**Figure 1L, M**) was sufficient to impair memory. With AAV-Syn-GFP used as a control vector, neither injection of AAV Syn-GFP into IL-4Rα^fl/fl^ mice nor injection of AAV Syn-cre virus into IL-4Rα^+/+^ mice altered freezing behavior (**Figure K, M**). These data suggest that neuronal expression of IL-4Rα is required for normal CFC memory processing.

### Rescue of synapse-related gene expression in CFC-activated neurons by T cells and IL-4

Peripheral immune cells have no physical contact with neurons in young-adult healthy mice, whereas their meningeal spaces are densely populated by a variety of immune cells (Kipnis, 2016; Mrdjen et al., 2018). We hypothesized that cytokines released into the cerebrospinal fluid (CSF) by meningeal immune cells diffuse through the brain parenchyma (Da Mesquita et al., 2018) and target their receptors on neurons and/or glia to influence memory. To determine how immune signals change neuronal function, we labeled WT, SCID and T-cell-repopulated SCID mice with AAV9-cFos-tTA and AAV9-TRE-H2B-GFP viruses in the hippocampal DG (**Figure 2A**, **Supplemental Figure S3A−F**) (Liu et al., 2012a). Labeled activated neuronal nuclei were isolated 24 hr after fear conditioning training, and snRNAseq was performed (**Figure 2B**), with non-activated neurons from the same brain region used as controls (**Figure 2C**). Analysis of neuronal nuclei from the DGs of WT, SCID and T-cell-repopulated SCID mice revealed unique clusters, distinctively falling into glutamatergic or GABAergic subsets (**Supplemental Figure S4A, B**). To gain an understanding of the molecular differences in task-activated neurons between WT and SCID mice we first examined the differential expression between the two groups, identifying 1,497 significant changes (**Figure 2D, E**). Among these changes, genes annotated to the Gene Ontology (GO) terms for synapse organization, regulation of ion transmembrane transport, and synaptic membrane adhesion were included in the top 10 of the most statistically significant observed (**Figure 2D, E**). Between WT and SCID groups, genes associated with synapse-related GO terms were also differentially expressed in non-activated neurons, suggesting that the adaptive immune system may regulate synapse-related gene expression under homeostatic conditions (**Supplemental Figure S4C−F**).

**Figure 2:**
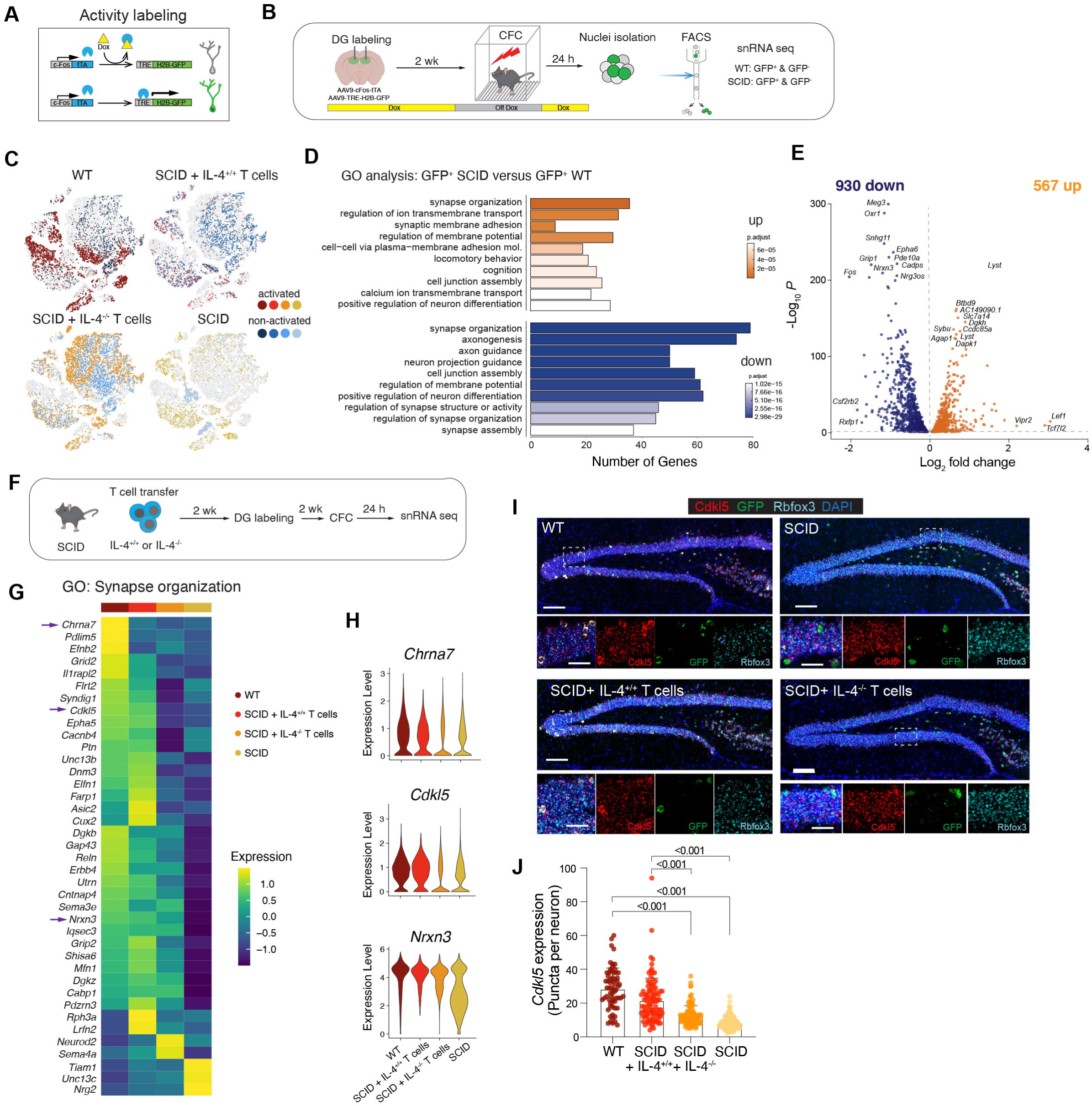
Transcriptomic changes in synapse-related genes are orchestrated by T cell-derived IL-4. **A,** Schematic representation of the experimental design. WT and SCID mice were injected with AAV9-cFos-tTA and AAV9-TRE-H2B-GFP and maintained on doxycycline (DOX) chow. **B**, The dentate gyrus was labeled two weeks prior to training. To specifically label training activated neurons, mice were taken off DOX food 24 hour before training and feed DOX chow immediately after training. Activated (GFP^+^) and non-activated (GFP^-^) neuronal nuclei from the DG were sorted by flow cytometry and subjected to single-nuclei RNA sequencing. **C**, t-SNE showing the distribution of cells from each sample (WT, SCID, SCID with IL4^+/+^ T cells, SCID with IL4^-/-^ T cells) of activated and non-activated neurons across clusters. **D**, Gene ontology analysis of top 10 pathways by significance comparing differentially expressed genes in activated GFP^+^ neuronal nuclei from SCID and WT mice. **E**, Volcano plot showing differentially expressed genes in activated WT compared to activated SCID neurons. Top differentially expressed genes are labelled with text. **F**, Schematic representation of the experimental design. SCID mice were injected with IL-4^+/+^ or IL-4^-/-^ splenic T cells and processed as described in A. **G,** Heat maps showing the average scaled expression of significantly differentially expressed genes in GO:0050808 (synapse organization) between WT, SCID with IL4^+/+^ T cells, SCID with IL-4^-/-^ T cells and SCID of activated neuronal nuclei. Mean of n=3-4 biological samples. **H**, Violin plots showing the distribution of expression of *Chrna7, Cdlk5, Nrxn3* in activated neuronal nuclei. **I**, Representative images of multi-color RNAscope experiments showing *Cdkl5, GFP* and *Rbfox3* gene expression of neurons in the dentate gyrus for conditions shown in **F.** Scale bars = 200 µm or 40 µm. **J**, Quantifications of *Cdkl5* gene expression in activated neurons (*GFP^+^ Rbfoxp^+^*) of WT (n = 64 neurons), SCID with IL4^+/+^ T cells (n = 95 neurons), SCID with IL-4^-/-^ T cells (n = 116 neurons) and SCID mice (n = 116 neurons). Data are presented as means ± SEM.; 3-4 mice per group. Two-way ANOVA with Bonferroni post hoc test.

We next determined those genes that are involved in the enriched “synapse organization” pathway and have been shown to play a vital role in excitability. Upon repopulation of SCID mice with T cells (**Figure 2F**), we identified the genes involved in positive regulation of synapse density. Interestingly, we found that repopulation of WT T cells in SCID mice largely reversed the differential gene expression between WT and SCID, both in CFC-activated neurons (**Figure 2G**) and in CFC-non-activated neurons (**Supplemental Figure S4E**). Violin plots of gene expression showed that in SCID mice, synapse-related genes such as *Chrna7* and *Cdkl5* were restored to WT levels by T-cell-repopulation with WT but not with IL-4^−/−^ T cell repopulation. Notably, expression of several genes such as *Nrxn3* were rescued by both WT T cell and IL-4^−/−^ T cell repopulation (**Figure 2H**). *Cdkl5*, in particular, has been reported to regulate synaptic function and memory (Ricciardi et al., 2012; Tang et al., 2017; Zhu et al., 2013). We therefore collected brains, 24 hr after exposure to foot shock, to further verify whether T cells and their derived IL-4 regulate *Cdkl5* expression in CFC-activated neurons. Using *in-situ* hybridization methods, we quantified *Cdkl5* expression on Rbfox3^+^-and GFP^+^-tagged neurons under four different conditions (**Figure 2I**). Consistently with our snRNAseq findings, RNAScope showed a significant increase in the number of activated *Cdlk5*^+^ neurons in the DG of mice reconstituted with T cells, compared to mice that remained without T cells (**Figure 2J**).

### IL-4 regulates CFC memory through IL-4R in GABAergic neurons

We next focused on identifying the specific neuronal subsets responsible for promoting fear memory following IL-4 signaling. In different types of neurons and brain regions in published data sets (**Supplemental Figure S5A−C**), we detected IL-4Rα mRNA in both excitatory and inhibitory neurons in the hippocampus (**Supplemental Figure S2A−C**) and further verified *Il4r* and *Il13r 1* expression (IL-4 signals through a heterodimer of IL-4Rα /IL13Rα1 or IL-4Rα common gamma chain (γ_c_)). While the percentages of cells expressing each receptor varied slightly across tissues and datasets, we confirmed that both receptors are expressed in both excitatory and inhibitory neurons, with particularly high expression of *Il4ra* in several inhibitory subpopulations (**Supplemental Figure S5B, C**). To determine the function of IL-4Rα in excitatory and inhibitory neurons for CFC memory regulation, we conditionally ablated IL-4Rα from either GABAergic (Gad2^Cre+^::IL-4Rα^fl/fl^) or glutamatergic (vGlut2^Cre+^::IL-4Rα^fl/fl^) neurons. Mice with IL-4Rα deficiency in GABAergic neurons (**Figure 3A**), but not in glutamatergic subsets (**Figure 3B**), exhibited significantly decreased freezing times, suggesting that in affecting CFC memory, IL-4 signals mainly through inhibitory neurons.

**Figure 3:**
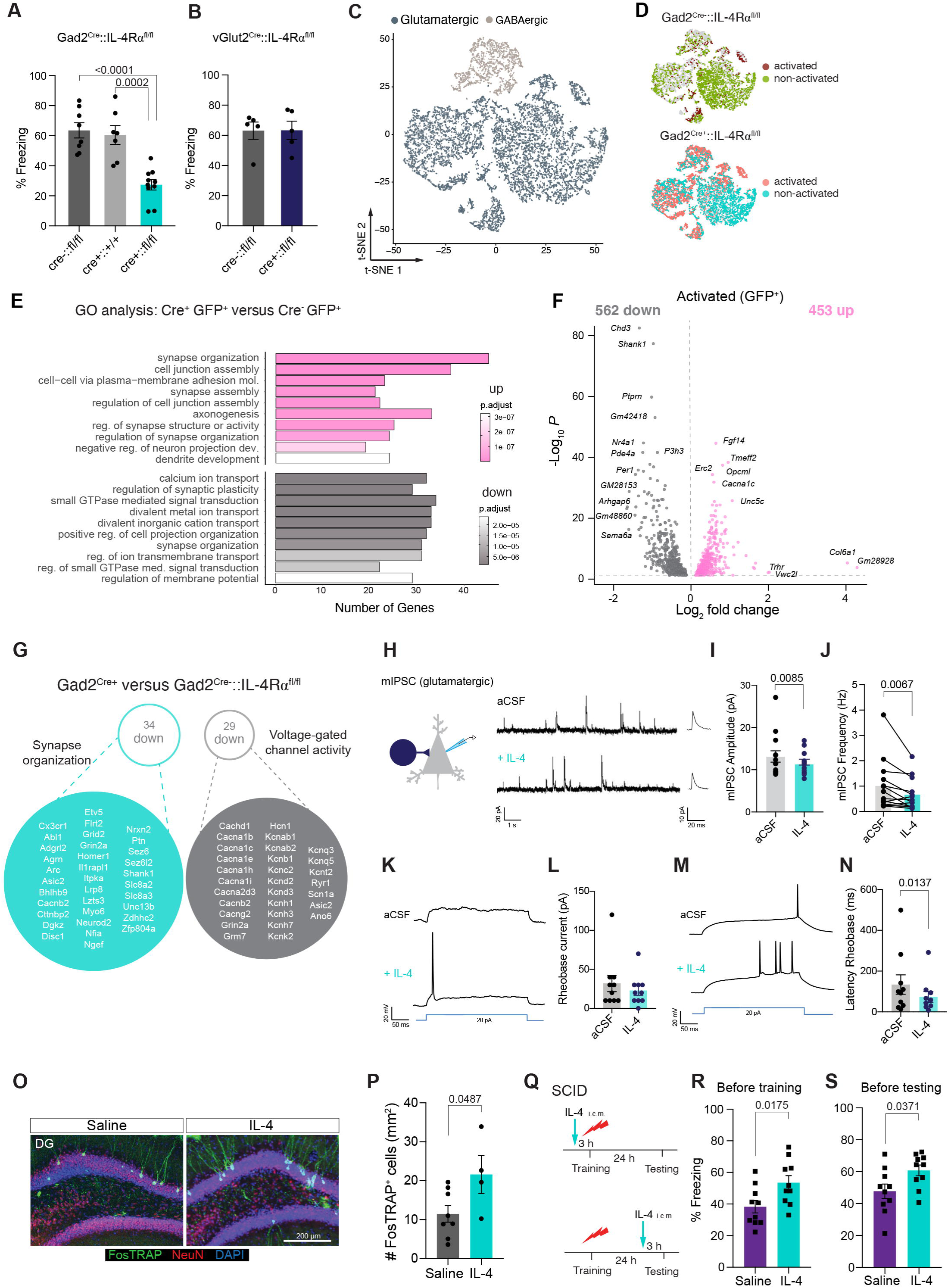
IL-4Rα on GABAergic neurons regulates contextual fear memory. **A,** Percent freezing time of mice with GABAergic neuron-specific knockout of IL-4Rα. Gad2^Cre-^ ::IL-4Rα^fl/fl^ (n= 8), Gad2^Cre+^::IL-4R^+/+^ (n= 7), and Gad2^Cre+^::IL-4Rα^fl/fl^ (n=10) mice. One-Way ANOVA with Tukey post hoc test. **B**, Percent freezing level of mice with glutamatergic neuron-specific knockout of IL-4Rα. vGlut2^Cre-^::IL-4Rα^fl/fl^ and vGlut2^Cre+^::IL-4Rα^fl/fl^ mice. N= 5 mice per group. Two-tailed unpaired Mann-Whitney *U*-test; no significant difference was found between the groups. **C**, t-SNE visualization of nuclei from the dentate gyrus of Gad2^Cre^::IL-4Rα^fl/fl^ mice, colored by glutamatergic and GABAergic neurons. **D,** t-SNE visualization of activated and non-activated nuclei in Gad2^Cre-^::IL-4R^fl/fl^ mice and Gad2^Cre+^::IL-4Rα^fl/fl^ mice. **E**, GO analysis of DEGs that were identified between activated GFP^+^ neurons from Gad2^Cre+^::IL-4R^fl/fl^ mice versus Gad2^Cre-^::IL-4Rα^fl/fl^ mice. **F**, Volcano plots showing differentially expressed genes in activated neurons in Gad2^Cre+^::IL-4Rα^fl/fl^ versus Gad2^Cre-^::IL-4Rα^fl/fl^ mice. Note, top 10 significantly altered genes (adjusted p < 0.05) with absolute log fold changes > 0.15 are annotated. **G**, List of synapse related and voltage-gated ion channels (calcium-, sodium-, potassium channels) that are differentially expressed in activated glutamatergic neurons from Gad2^Cre+^::IL-4Rα^fl/fl^ mice and Gad2^Cre-^::IL-4Rα^fl/fl^ mice based on GO:0050808, GO:0022843 and GO:0022844. **H**, Representative electrophysiological recording showing miniature inhibitory post synaptic current (mIPSC) in DG excitatory neurons before and after perfusion with 100 ng/ml IL-4. **I-J**, Statistical analysis of changes in mIPSC amplitude (**I**) and frequency (**J**) before and after IL-4 perfusion (n= 14 neurons from 9 mice). Two-tailed paired Wilcoxon test. **K**, Examples showing DG excitatory neuronal action potential response upon injection of current during rheobase (the minimum current injection required for action potential) test before and after IL-4 perfusion. Two-tailed paired Wilcoxon test. **L**, Statistical analysis of rheobase current before and after IL-4 perfusion (n= 10 neurons/group). **M**, Examples showing latency of the response to rheobase in glutamatergic neurons in DG region before and after IL-4 perfusion. **N**, Statistical analysis of latency of the response to rheobase in glutamatergic neurons in the DG before and after IL-4 perfusion. (n= 10 neurons/group). Two-tailed paired Wilcoxon test; Each dot indicates one individual neuron. Data are presented as the mean SEM. **O**, Representative micrographs showing neuronal labeling in the dentate gyrus (DG) with cFosTRAP mice subjected to saline or IL-4 i.c.m. injection. Scale bar = 200 µm. **P,** Quantifications of FosTRAP^+^ neurons upon IL-4 injection. **Q**, Experimental schematics showing SCID mice either injected with saline or 100 units of IL-4 (IL-4) through the intra-cisterna magna (i.c.m.) 3 hours before training. Two-tailed unpaired Students *t*-test. **R** - **S**, Percent freezing time of SCID mice injected with saline or IL-4, 3 hours before training (**R**) or testing (**S**). Two-tailed unpaired Mann-Whitney *U*-test.

Because our data on IL-4Rα depletion pointed to involvement of GABAergic neurons in CFC memory regulation, we conducted snRNAseq experiments on CFC-activated neurons from the DG using Gad2^Cre–^::IL-4Rα^fl/fl^ and Gad2^Cre+^::IL-4Rα^fl/fl^ mice (**Figure 3C−G**). Most strikingly, loss of IL-4Rα in Gad2^+^ (inhibitory) neurons led to changes in expression of genes of relevance to GO terms, such as “synapse organization”, “regulation of synaptic plasticity”, and “regulation of synapse structure or activity” (**Figure 3E**), indicating that IL-4 may regulate synaptic function through IL-4Rα on inhibitory neurons. Moreover, loss of IL-4Rα in Gad2^+^ (inhibitory) neurons also led to lower expression of genes related to voltage-gated potassium, sodium, and calcium channels in glutamatergic neurons in the DG (**Figure 3G**), suggesting that IL-4 may also regulate neuronal excitability.

To determine whether IL-4Rα signaling can drive changes in neuronal activity in different types of neurons, we recorded AMPA-type glutamate-receptor-mediated miniature excitatory postsynaptic currents (mEPSC) and GABAA receptor-mediated miniature inhibitory postsynaptic currents (mIPSC) in excitatory and inhibitory neurons in the DGs of acutely prepared brain slices from WT mice. IL-4 perfusion significantly decreased mIPSC frequency and amplitude in excitatory neurons (**Figure 3H−J**). In excitatory neurons, IL-4 perfusion also did not change mEPSC frequency or amplitude (**Supplemental Figure S6B−D**), or mIPSC and mEPSC frequencies or amplitude in inhibitory neurons (**Supplemental Figure S6E−J**). Lastly, IL-4 perfusion was found to decrease the latency of the minimum injection of current required for an action potential (rheobase) in glutamatergic neurons, suggesting that IL-4 can increase glutamatergic neuronal excitability (**Figure 3K−N**).

### IL-4 regulates synaptic transmission and restored memory deficits in SCID mice

Finally, we focused on the functional consequences of the IL-4 effects on neurons, evaluating the isolated effects of IL-4 because of its profound positive outcomes in SCID mice after T-cell transfer. To enable activated neurons to be labeled *in vivo* and during a specific time window (in the presence of 4-hydroxytamoxifen (4-OHT)), we used mice that express Cre^ER^ under the control of an activity-dependent c-Fos promoter (FosTRAP mice (Guenthner et al., 2013)) crossed with Ai6 mice (Rosa-CAG-LSL-ZsGreen1) to capture neurons that were active during IL-4 administration. Relative to saline-injected controls, injection of IL-4 into the CSF significantly increased the number of activated FosTRAP^+^ neurons in the DG (**Figure 3O, P**).

We next examined whether IL-4 plays a role in the behavioral reversal of a deficit in CFC memory. Remarkably, a single injection of recombinant IL-4 into the CSF of SCID mice 3 hr before CFC training improved memory deficits (**Figure 3Q, R**). Administration of IL-4 following task acquisition and 3 hr before memory retrieval also led to an increased freezing time (**Figure 3Q, S**). However, freezing behavior was not changed by injection of recombinant IL-4 into the CSF of WT mice before either training or testing (**Supplemental Figure S7A, B, C**), suggesting that the IL-4-mediated effects on memory are homeostatic, i.e., that the process is exacerbated in absence of IL-4, but is not affected by boosting it beyond physiological levels. Collectively, these data demonstrate that interference with CD4 T cells and their derived IL-4-mediated functions fosters transcriptional dysregulation, synaptic weakening in neurons, and an inability to fully operate during fear memory.

## Discussion

Appropriate defensive responses to previous experiences are vital in promoting survival when the individual is confronted with a potential threat. By studying an episodic memory task of CFC in mice, we showed here that T-cell-derived IL-4 plays a key role in regulating this type of memory. Mice depleted of CD4 T cells, or lacking adaptive immunity (SCID), as well as deficient in IL-4, exhibit memory deficits. In SCID mice, memory could be restored by reconstitution with T cells from WT donors but not with T cells from donors deficient in IL-4. Memory in SCID mice could also be improved by injection of recombinant IL-4 into the CSF either before acquisition of the CFC task or before memory retrieval. While memory function in the mice was not affected by conditional deletion of IL-4Rα from their microglia, deletion of IL-4Rα from their neurons was sufficient to recapitulate the cognitive deficits seen in IL-4-deficient mice. *In-situ* hybridization and snRNAseq data showed that IL-4Rα is expressed in both excitatory and inhibitory neurons, yet deletion of IL-4Rα from GABAergic neurons was required to recapitulate memory deficits. By comparing transcriptomic profiles of task-activated neurons in T-cell-sufficient and T-cell-deficient mice, we found that many of the differentially expressed genes were related to synaptic function.

The inhibitory activity of GABAergic neurons was increased in the presence of IL-4, and their synaptic properties were impaired in the absence of IL-4 receptor. snRNAseq of task-activated neurons from mice with specific IL-4Rα knockout in GABAergic neurons and their WT littermates showed that removal of IL-4Rα from GABAergic neurons alters the expression of genes associated with synapse organization and synaptic plasticity. This suggested that IL-4 could be an important contributory force in synapse reorganization, leading to strengthening of inter-neuron connections and improved cognition.

The results presented here support the existence of a mechanistic link between adaptive immunity and neuronal function (Ben-Shaanan et al., 2016; Ben-Shaanan et al., 2018; Filiano et al., 2016b), and extend those interactions to physiological homeostatic conditions. Our findings demonstrate a direct effect of peripheral T cells and their derived IL-4 on behavior modulation, neuronal transcription and synaptic function in CFC.

Many of the “immune” genes and pathways that are upregulated in activated neurons have previously been linked to neuronal function. As examples, TNF signaling was implicated in synaptic scaling (Stellwagen and Malenka, 2006), complement molecules were shown to modify synaptic function (Stevens et al., 2007), and MHCI was found to be expressed by neurons and to directly regulate synaptic pruning and function (Shatz, 2009). It is therefore plausible that “classical” immune molecules such as IL-4, which are expressed primarily by immune cells, can be used by the immune system for signaling to neurons about the state of the organism. Such signaling would be performed directly on neurons expressing receptors for such immune molecules. Cytokine signaling would then modulate neuronal function, in part through “shared” immune molecules (such as TNF, MHCI and complement system-related molecules, all of which are expressed by neurons). Future studies should focus on cytokine-induced subcellular signaling to neurons and, more specifically, on the role of such shared immune molecules in mediating neuro-immune communications. Thus, for example, while it is likely that T cells affecting brain function are harbored in the meninges (Radjavi et al., 2014), in order to further address this communication enigma there is a critical need for new tools that would allow spatial targeting of T cells within such compartments and in tissues of interest.

Since the immune system is more easily accessible than the brain, modification of immune cells may indeed turn out to be a promising therapeutic option for disorders that are not classically characterized as neuroinflammatory, but in which an immune link is present.

## Materials and Methods

### Mouse strains and housing

Wild-type mice (C57BL/6J background) were either bred in-house or purchased from the Jackson Laboratory (JAX 000664). All mice were maintained in the animal facility for at least one week prior to the start of any experiment. The following strains were used: C57BL/6-*Il4^tm1Nnt^*/J (IL-4^−/−^, JAX 002518), B6.CB17-*Prkdc^scid^/*SzJ (SCID, JAX 001913), B6J.Cg-*Gad2^tm2(cre)Zjh^*/MwarJ (Gad2-IRES-Cre, JAX 028867), B6J.129S6(FVB)-*Slc17a6^tm2(cre)Lowl^/*MwarJ (Vglut2-IRES-Cre, JAX 028863), B6.129(Cg)-*Fos^tm1.1(cre/ERT2)Luo^*/J (*Fos^CreEr^*, JAX 021882), B6.Cg-*Gt(ROSA)^26Sortm6(CAG-ZsGreen1)Hze^*/J (Ai6, JAX 007906), B6;129S-Gad2tm1.1Ksvo/J (Gad2-T2a-NLS-mCherry, JAX 023140), B6.Cg-Tg(Syn1-cre)671Jxm/J (Syn-cre, JAX 003966) and B6.129P2(Cg)-*Cx3cr1^tm2.1(cre/ERT2)Litt^*/WganJ (Cx3cr1^CreERT2^, JAX 021160). The *Il4ra*^tm2Fbb^ (IL-4Rα^fl/fl^) mice were kindly provided by Frank Brombacher (University of Cape Town). All mice were housed under standard 12LJ light:dark cycle conditions (lights on at 7:00am) in rooms with controlled temperature and humidity. They were given standard rodent chow and sterilized tap water *ad libitum* unless stated otherwise. Male mice at 10 to 20 weeks of age were used for behavior experiments unless stated otherwise. All experiments were approved by the Institutional Animal Care and Use Committee of the University of Virginia.

### Contextual fear conditioning (CFC) training

The equipment was purchased from Coulbourn Instruments. During training, the test cage was scented with 0.25% benzaldehyde. Each mouse was put in an isolation cubicle test cage. After 3 min of habituation in the test cage (context B), electric foot-shocks (2 sec, 0.50 mA) were delivered during 198-200 sec, 258-260 sec, 318-320 sec. Mice were kept in the test cage for additional 30 sec after training and were put back to their home cage. Animal groups were blinded to the observer and video images analyzed offline and manually scored. Freezing was defined as an event where the entire mouse was sitting still.

### Intra-cisterna magna injection of IL-4

Mice were anesthetized with 1.5% isoflurane. The skin of the neck shaved and sterilized with iodine. The head of the mouse was gently secured in a stereotaxic frame. A skin incision was made, and the muscle layers were retracted to expose the cisterna magna. Using a Hamilton syringe with a 33-gauge needle, 3 μl of 100 U of murine IL-4 (eBioscience, 14-8041-80) or saline was injected into the cisterna magna compartment. The syringe was left in place for an additional 2 minutes to minimize backflow of CSF. The skin was sutured, and the mice were allowed to recover for at least 3 hours before any behavioral experiments.

### T cell depletion

Mice were injected i.p. with 200 ug anti-CD4 (GK1.5, BioXCell) or isotype control antibody (LTF-2, rat IgG2b, BioXCell). To ensure continuous depletion, mice were injected i.p. with 100 ug weekly.

### TRAP labelling of activated cells

*Fos^CreER^* mice were crossed to *Ai6* mice to generate double heterozygous (*Fos^CreER^*::*Ai6*) mice used for the labelling experiments. The mice were injected in the intra-cisterna magna (i.c.m.) compartment with 100 U of murine IL-4 (eBioscience, 14-8041-80) diluted in saline at a total volume of 3 μl. As control, mice were injected with the same volume of saline. After two hours, the mice were given an intraperitoneal injection of 10 mg kg^-1^ of 4-hydroxytamoxifen (4-OHT; Sigma, H6278) dissolved in a 1:4 mixture of castor oil (Sigma, 259853): sunflower oil (Sigma, S5007). The drug preparation has been previously described (Guenthner et al., 2013). After one week, mice were sacrificed, and the brains were harvested for further analysis.

### Seizure induction

Mice were injected with 25 mg per kg kainic acid (Sigma-Aldrich, 420318) intraperitoneally and seizure behavior was monitored. Animals progressed at least to stage 3 on a Racine scale (continuous head bobbing and forepaws shaking) and were scarified 1 d after seizure induction.

### Construction of the pAAV-TRE-H2B-GFP plasmid

The University of Massachusetts Viral Vector core constructed the pAAV-TRE-H2B-GFP plasmid. The H2B-GFP fragment was amplified from the CMV-H2B-GFP (Addgene plasmid #11680) plasmid using the following primers: H2B-GFP-Forward: 5’-*gctgcggaattgtacccgcggccgatccaccggtcgccaccatggatgccagagcc agcg*-3’ and H2B-GFP-Reverse: 5’-*gcacagtcgaggctgatcagcgagctctagtcgacggtatcgatttacttgtacagctcgtccat gccgagagtga*-3’. The amplified fragment was cloned into the pAAV-TRE-EGFP backbone after restriction enzyme digestion with *Nco*I and *Cla*I to replace the original GFP sequence with H2B-GFP instead.

### Preparation of adeno-associated virus constructs

The following plasmids were purchased from Addgene: pAAV-cFos-tTA (Addgene plasmid #66794), and CMV-H2B-GFP (Plasmid #11680). The plasmids were packaged with AAV9 coat proteins in the Viral Vector Core of the Gene Therapy Center at the University of Massachusetts Medical School. AAV9-TRE-H2B-GFP was constructed by the University of Massachusetts Viral Vector core. Viral titers were 2×10^13^ genome copies (GC) mL^-1^ for AAV9-cFos-tTA, 1.5×10^13^ GC mL^-1^ for AAV9-TRE-H2B-GFP.

### Immunohistochemistry

Mice were given a lethal dose of anesthesia by intraperitoneal (i.p.) injection of euthasol (10% v/v in saline). They were transcardially perfused with ice-cold PBS with heparin (10 U ml^-1^) followed by 4% paraformaldehyde (PFA). Brains were dissected and kept in 4% PFA overnight at 4°C. The fixed brains were washed with PBS, cryoprotected in 30% sucrose at 4°C until sunk to the bottom of the vial, and frozen in Tissue-Plus OCT compound (Thermo Scientific). Frozen brains were sliced into 40μm-thick free-floating coronal sections using a cryostat (Leica). Brain sections were blocked with 1% bovine serum albumin (BSA), 2% normal serum (either goat or chicken), and 0.2% Triton X-100in PBS for 1h at room temperature (RT). The blocking step was followed by incubation with rabbit anti-NeuN-Alexa Fluor 647 or 488 (1:500, Abcam, ab190565 or ab190195, clone EPR12763) in PBS containing 1% BSA and 0.2% Triton X-100 overnight at 4°C. Sections were washed two times for 15 min at room temperature (RT) with PBS and nuclei were stained with 4’, 6-Diamidino-2-phenylindole (DAPI, 1:10,000, Sigma-Aldrich) at RT for 15 min. The tissues were washed with PBS once and were mounted with Prolong Gold (ThermoScientific).

### RNAscope® multiplex fluorescent assay

Euthanized mice were perfused with PBS containing heparin (10 U ml^-1^). Within 5 min, brains were embedded in Tissue-Plus OCT compound and were immediately frozen on dry ice. Frozen brains were sectioned into 16μm-thick coronal slices, mounted onto Superfrost plus slides and pretreated according to the manufacturer’s instructions for fresh frozen sample (RNAscope® Multiplex Fluorescent Reagent Kit v2 Assay). Briefly, sections were fixed in pre-chilled 10% neutral buffered formalin (Fisher Scientific) for 15 minutes at 4°C and were dehydrated using an ethanol series (50%, 70%, and 100% ethanol). The sections were pretreated with hydrogen peroxide for 10 min at RT and incubated with protease IV for 30 min at RT, provided in the kit. The hybridization and amplification steps were performed using *Il4ra* (Mm-IL4ra in C1), *Gad1* (Mm-Gad1-C2), *Gad2* (Mm-Gad2-C3), *Cdkl5* (Mm-Cdkl5-C2), *Rbfox3* (Mm-Cdkl5-C3), *Vglut1* (Mm-Slc17a7-C3) and *Vglut2* (Mm-Slc17a6-C2) RNAscope probes according to manufacturer’s instructions. Three-plex detection of IL-4Ra on glutaminergic and GABAergic neurons was achieved using Opal 520 (1:1000, Perkin Elmer), Opal 570 (1:1500, Perkin Elmer), and Opal 690 (1:1000, Perkin Elmer). Nuclei were stained with DAPI at RT for 30 sec and sections mounted with ProLong Gold antifade reagent (Invitrogen) were placed on the slide and covered with coverslips.

RNAscope® multiplex fluorescent images of the DG, with a 20× objective with 0.70 NA. Quantitative analysis was performed using Fiji by a blinded experimenter. Nuclei were segmented automatically in the DAPI channel and subsequently measured in each channel. For multiplex fluorescent image analysis, we used thresholding, watershed segmentation to identify and also split overlapping nuclei. Adjust the size and circularity to count cells only and avoid counting fragments. Visually inspect all the detected particles and exclude the particles that do not look like a real signal. Expend the area of identified nuclei to a proportion and measure the puncta for target channel. For multiple channel measurement, repeat this method. In R, nuclear events were refined using size and DAPI mean fluorescent intensity, and cell type markers (Gad1, Gad2, Vglut1, Vglut2) were plotted against IL-4Ra using ggplot2.

### Stereotactic injections

Mice were anaesthetized by i.p. injection of ketamine (100 mg kg^-1^) and xylazine (10 mg kg^-1^) in saline solution. The head of the mouse was secured in a stereotaxic instrument with a digital display console (KOPF, Model 940). Viruses were injected using a glass micropipette (WPI, 1B120F-4) filled with mineral oil attached to Nanoliter Injector with SMARTouch Controller (WPI, NL2010MC2T). The needle was slowly lowered to the target brain region and was left in place for 5 min before starting the injection. The injection speed was set at 40-50 nl min^-1^. After injection, the needle was left in place for an additional 10 min to minimize backflow. The mice were then injected subcutaneously with ketoprofen (2 mg kg^-1^) and allowed to recover on a heat pad until fully awake.

### Activity labeling of neurons

Mice were put on doxycycline food (ENVIGO, 40 mg kg^-1^) for 1 week. The mice were injected bilaterally with 300 nl of a 1:1 mixture of AAV9-cFos-tTA and AAV9-TRE-H2B-GFP into the retrosplenial cortex (AP: −2.06 mm; ML: ±0.4 mm; DV: 1.0 mm) or dentate gyrus (AP: −2.06 mm; ML: ±1.3 mm; DV: 1.9 mm). Two weeks after virus injection, mice were taken off Dox and were given standard rodent chow for 24 hours before the CFC training. After training, the mice were given doxycycline food. Twenty-four hours after training, mice were prepared for single nuclei sequencing.

### Long-term potentiation recordings

After deeply anesthetized with isoflurane (Henry Schein, USA), mouse was quickly decapitated, and the head was dipped into ice-cold artificial cerebrospinal fluid (aCSF) briefly. The brain was taken out by swift dissection and submerged into ice-cold aCSF that was bubbled continuously with 5% CO2/95% O2. The aCSF was constituted as (in mM) NaCl 124.0, KCl 2.5, KH2PO4 1.2, CaCl2 2.4, MgSO4 1.3, NaHCO3 26.0 and glucose 10.0 (pH 7.4). The right hippocampus was taken out and cut into slices (400 µm thick) along its longitude axis with posterior part first using a tissue slicer (Stoelting Co., IL, USA). The left hippocampus was cut into slices in a similar manner but was never used in this study. After at least 1 hour of incubation at room temperature in aCSF bubbled continuously with 5% CO2/95% O2, slices were recorded in submerged mode on the top of a chamber (Harvard Apparatus, USA) at room temperature with illumination underneath (Leica, Germany). Data were collected with an Axopatch-2B amplifier and pClamp 10.4 program (Molecular Devices, US) through Digidata 1320A. Slices were perfused continuously with aCSF bubbling with 5% CO2/95% O2 at flow rate about 2 ml/min using two channels peristaltic pump (Dynamax, Rainin, USA).

The recording electrode (resistance 1-3 MΩ) was was fabricated with capillary glass tubing (Warner Instruments, Model No: G150F-3) and filled with aCSF. Biphasic current pulses (0.2 ms duration for one phase, 0.4 ms in total) were delivered in 10 s intervals using a concentric bipolar stimulating electrode (CBARC75, FHC, USA) through an isolator (ISO-Flex, AMPI, Israel). To record field population spikes (PS) in dorsal dentate gyrus (DG), the recording electrode was placed on inner part of granular cell layer and the stimulating electrode was placed right above hippocampal fissure to stimulate the bypassing perforant pathway fibers. To record field excitatory postsynaptic potential (EPSP) in DG the recording electrode was placed on the middle stratum above granular cell layer and stimulating electrode was placed above the hippocampal fissure. Input/output curve was obtained for each slice by increasing intensities of 1 threshold (T, typically 0.02mA in CA1 and 0.05 mA in DG). The maximum intensity did not pass 1mA. For baseline recording, test pulse intensities were adjusted to evoke EPSP about 35-50% of maximal response. Theta burst (12 trains of 4 pulses at 100 Hz, delivered at 5 Hz, repeated three times at 0.1 Hz) and the amplitude of population spike and slope of EPSP were measured from the initial phase of negative wave with pClamp 10.4. Each data point was the average of 3 consecutive traces. LTP was plotted as the percentage to 10 min recording of baseline following high frequency stimulation. Representative trace is the average of 6 consecutive ones which last for 1 minute in span. Data analysis was processed with Excel (Microsoft, USA). The last 5 min of LTP level values were averaged for comparison among groups. Data were presented as mean ± Sem.

### mIPSC and mEPSC patch-clamp recordings

Mice (wild type C57BL/6J or B6.129S-Gad2^tm1.1Ksvo^/J, specified otherwise, both sexes, at 10 - 22 postnatal days old, P10–P22), were decapitated under anesthesia with isoflurane (USP, Henry Schein). The animal’s brain was rapidly dissected and transferred to ice-cold (∼4°C) oxygenated sucrose-substituted artificial cerebrospinal fluid (aCSF) solution. Coronal brain slices 250-μm thick were prepared with a Vibratome (Leica VT1000S) in oxygenated cold sucrose-substituted aCSF. The slices were transferred into an incubation chamber with normal aCSF, composing of (in mM) 130 NaCl, 1.2 NaH2PO4, 2.4 CaCl2, 2.5KCl, 1.3 MgSO4, 26 NaHCO3, 10 dextrose (pH 7.30, ∼300 mOsm, oxygenated at 95% O2/5% CO2), for at least 1h at room temperature (25 ± 2 °C) prior to electrophysiological studies.

For patch clamp recordings, individual slices were transferred to a submersion recording chamber and were continuously perfused with 95% O2/5% CO2 bubbled with the aCSF. Recordings were made from neurons in dentate gyrus using a MultiClamp 700B amplifier (Molecular Devices, Sunnyvale, CA, USA). mIPSC or mEPSC recordings of hippocampal dentate gyrus neurons were performed for 10 min as baseline. The chamber was perfused with 100 ng/ml recombinant IL-4 (eBioscience, 14-8041-80) and the same neurons recorded for another 10 min. Additional 10 min were recorded under continuous perfusion with aCSF and are indicated as washout. Signals were low-pass filtered at 1 kHz and sampled at 10 kHz using Digidata 1440A (Molecular Devices). For mEPSC recordings, glass pipettes (resistance, 3–5 mΩ) were loaded with internal solution containing (in mM): 110 K-gluconate, 20 KCl, 20 HEPES, 5 MgCl2, 0.6 EGTA, 2 MgATP, 0.2 Na3GTP (pH 7.3, ∼290 mOsm), and all neurons were held at −60 mV in voltage-clamp mode; 10 µM bicuculline methiodide and 1 µM tetrodotoxin (TTX) were added to aCSF to block GABAA receptor-mediated and voltage-gated Na^+^ currents, respectively. For mIPSC recordings, internal solution containing (in mM): 140 Cs-gluconate, 1 MgCl2, 11 EGTA, 1CaCl2 and 10 HEPES (pH 7.3, ∼290 mOsm) was used, and cells were held at 0 mV in voltage-clamp mode; NBQX (30µM) and TTX (1 µM) were added to block AMPA receptor-mediated and Na^+^ currents, respectively. The spike frequency versus injected current experiments were performed in current-clamp mode. The internal solution contained (in mM): 110 K-gluconate, 20 KCl, 20 HEPES, 5 MgCl2, 0.6 EGTA, 2 MgATP, 0.2 Na3GTP (pH 7.3, 290 mOsm). The excitatory and inhibitory synaptic blockers (50 µM D-APV, 30 µM NBQX, and 10 µM bicuculline methiodide) were added to the aCSF. The action potential firing rate was averaged from 10 depolarizing current injections (500 ms in duration at an interval of 2 s). The maximum current injected in each experiment was below the current that induced spike frequency adaptation.

### Adoptive transfer

Spleens, inguinal and cervical lymph nodes were collected. They were homogenized, and the cell suspension was passed through a 70-um cell strainer (Fisher Scientific). The red blood cells were lysed using ammonium-chloride-potassium (ACP) lysis buffer (Quality Biological) for 4 min at RT. The cells were washed once with RPMI (Gibco) supplemented with 5% fetal bovine serum (FBS). The single cell suspension was counted, pelleted, and adjusted so that the cell concentration was 1×10^8^ cells per ml in sterile saline. At 4 weeks of age, SCID mice were reconstituted with 5×10^6^ splenocytes or were given saline by intravenous (i.v.) injection. For T cell transfer, T cells were isolated from single-cell suspension from wild-type and IL-4^−/−^ mouse spleens using the Pan T cell Isolation Kit II, LS columns, and a QuadroMACS Separator (Miltenyi Biotech) for manual cell enrichment. Purified T cells were resuspended in saline, and 2×10^6^ cells (at a final volume of 100 μL) or sterile saline were injected into SCID recipients.

### Flow cytometry

Mice were given a lethal dose of anesthesia by intraperitoneal (i.p.) injection of Euthasol (10% v/v in saline) and were transcardially perfused with ice-cold PBS containing 10 U mL^-1^ heparin. The spleen and dural meninges were dissected and digested in 1 mg mL^-1^ collagenase 8, D and 5 U mL^-1^ DNase I (Sigma Aldrich) at 37°C for 30 min. The digestion was stopped by adding EDTA to a final concentration of 10 mM. The cell suspension was passed through a 70-um cell strainer. Cells were washed with RPMI (Gibco), resuspended in PBS, and counted with a cell counter (Nexcelom). Cells were stained with Zombie Aqua™ dye (Biolegend) for 20 min on ice. Cells were washed with PBS containing 2% FBS (FACS buffer), and Fc-receptors were blocked with anti-CD16/32 antibody in 50 μl total volume. An equal volume of primary antibody mix was added containing anti-CD45.2-Alexa Fluor 700, anti-CD4-eFluor450, anti-CD8a-PE, anti-CD90.2-PE-Cy7, and anti-TCRβ-FITC. Cells were stained for 30 min on ice and washed twice with FACS buffer. The data was acquired on a flow cytometer (Gallios Beckman Coulter) and analyzed using FlowJo software (Tree Star, Inc.) or Cytobank Premium.

### Single-nuclei RNA sequencing

Brains from mice injected with AAV9-cFos-tTA and AAV9-TRE-H2B-GFP in the retrosplenial cortex or dentate gyrus were rapidly extracted after perfusion with ice-cold artificial cerebrospinal fluid (aCSF). The brains were sectioned in aCSF with a vibratome (Leica) to obtain 500 um-thick slices. The retrosplenial cortex or dentate gyrus was micro-dissected, and tissues were pooled from 3-4 mice of the same group and experimental condition. The sections were then placed into working nuclei isolation medium (NIM; 0.25 M sucrose, 25 mM potassium, 5 mM magnesium chloride, 10 mM TrisCl, 100 mM dithiothreitol, 1x protease inhibitor, RNase inhibitor) and triturated with a 1000 μl wide-bore pipette tip. After adding 0.1% Triton-X-100, samples were homogenized using a 2 ml Dounce homogenizer with pestle A, followed by pestle B (Sigma), and filtered through a 70-um cell strainer (Fisher Scientific). The homogenate was spun at 1000 g for 8 min at 4°C, pellets, washed once, and resuspended in 500 μl 1% BSA in PBS for staining. Dissociated nuclei were stained by adding Hoechst 33342 (Invitrogen, 1:1000) and anti-NeuN-Alexa Fluor 647 antibody (Abcam, 1:500, ab190565, clone EPR12763) for 15 min on ice. Samples were then washed and resuspended in 1% BSA in PBS for FACS sorting on a BD Influx sorter. Single neuronal engram (Hoechst^+^ NeuN^+^ GFP^+^) or non-engram (Hoechst^+^ NeuN^+^ GFP^−^) nuclei were directly deposited into 1.5 ml tubes containing 0.04% non-acetylated BSA in PBS, and diluted to 1000 nuclei per ul estimated from counting on a hemocytometer and Trypan blue staining. The sorted cortical neuronal nuclei (∼2000) per sample were loaded onto a 10X Genomics Chromium platform to generate cDNAs carrying cell- and transcript-specific barcodes and sequencing libraries constructed using the Chromium Single Cell 3’ Library & Gel Bead Kit 2. Libraries were sequenced on the Illumina NextSeq using paired-end sequencing, resulting in 100,000 reads per cell.

### WT and SCID with IL-4^+/+^ or IL-4^-/-^ splenic T cell transfer single-nuclei analysis

#### Data Pre-processing

Base call files were converted to Cellranger compatible Fastq files using the Illumina Bcl2fastq2 software. A reference pre-mRNA genome was created using the *mkref* functionality in the Cellranger software using the mm10 genome Fastq file and an edited GTF file with the feature type for each transcript set to exon to bypass the Cellranger exon filtering and to include intronic counts for further analysis. Reads were then aligned to the mm10 pre-mRNA genome using the *count* function of the Cellranger software pipeline (version 3.0.2) provided by 10x genomics. Base call files were converted to Cellranger compatible Fastq files using the Illumina Bcl2fastq2 software. A reference pre-mRNA genome was created using the *mkref* functionality in the Cellranger software using the mm10 genome Fastq file and an edited GTF file with the feature type for each transcript set to exon to bypass the Cellranger exon filtering and to include intronic counts for further analysis. Reads were then aligned to the mm10 pre-mRNA genome using the *count* function of the Cellranger software pipeline (version 3.0.2) provided by 10x genomics. The resulting filtered gene by cell matrices of UMI counts for each sample were read into R using the read10xCounts function from the Droplet Utils package. Filtering was applied in order to remove low quality cells, excluding those with reads fewer than 2,000 or greater than 60,000, or unique genes fewer than 500 or greater than 8,000, or greater than 15% mitochondrial gene expression. Expression values for the remaining cells were then normalized using the scran and scater packages and the resulting log2 values were transformed to the natural log scale for compatibility with the Seurat (v3) pipeline (Butler et al., 2018; Lun et al., 2016; McCarthy et al., 2017).

#### Dimensionality Reduction and Clustering

The filtered and normalized matrix was used as input to the Seurat pipeline and cells were scaled across each gene before the selection of the top 2,000 most highly variable genes using variance stabilizing transformation. Principal Components Analysis was conducted, and an elbow plot was used to select the first six principal components for tSNE analysis and clustering. Shared Nearest Neighbor (SNN) clustering optimized with the Louvain algorithm, as implemented by the Seurat *FindClusters* function was performed before manual annotation of clusters based on expression of canonical gene markers. Cell types not of interest to the analysis, namely glia, were removed and the remaining neuronal clusters were re-scaled, re-clustered, and identified as above. Clusters were then collapsed based on common cell types to result in two clusters.

#### Differential Expression

For analysis of differentially expressed genes between conditions, Glutamatergic neurons were isolated and filtered to include genes that had at least 5 transcripts in at least 5 cells, then the top 2000 highly variable genes were determined and included for further analysis using the SingleCellExperiment *modelGeneVar* and *getTopHVGs* functions. After filtering, observational weights for each gene were calculated using the ZINB-WaVE *zinbFit* and *zinbwave* functions (Berge et al., 2018). These were then included in the edgeR model, which was created with the *glmFit* function, by using the *glmWeightedF* function (Robinson et al., 2009). Results were then filtered using a Benjamini-Hochberg adjusted p-value threshold of less than 0.05 as statistically significant. Volcano plots were created with the EnhancedVolcano package in R (Blighe et al., 2020).

#### Pathway Enrichment

Over representation enrichment analysis with Fisher’s Exact test was used to determine significantly enriched Gene Ontology (GO) terms (adj. p < 0.05) for the sets of significantly differentially expressed genes between conditions. For each gene set, genes were separated into up- and down-regulated and separately (Hong et al., 2013) the *enrichGO* function from the clusterProfiler package was used with a gene set size set between 10 and 500 genes and p-values adjusted using the Benjamini-Hochberg correction (Yu et al., 2012, 2015).

### Analysis of Publicly Available Datasets

The publicly available RiboTag data was downloaded from the GEO portal (GSE133291) and expression levels of genes of interest were replotted using ggplot2 (PMID:31451803). Publicly available count data (GSE92522) was downloaded from the Gene Expression Omnibus website and reanalyzed in R using the scran and scater packages for normalization and the Seurat workflow for further analysis including scaling, principal component analysis, clustering and tSNE. Violin plots were generated using the Seurat package and the percentage of *Il4ra* expression was calculated within each of the subpopulations identified in the provided metadata sample guide (PMID:2894923).

The level 6, taxonomy level 2 dataset for all CNS neurons was downloaded from the mousebrain.org website in loom format and analyzed in python to extract count information for cells isolated from the Dentate Gyrus, CA1 hippocampus (CA1), Hippocampus (HC), Cortex (Ctx1, Ctx1.5, Ctx2, and Ctx3), the SS Cortex, Amygdala, Thalamus, and Hypothalamus as well as information about the Linnarsson identified cluster to which each cell belongs (Zeisel et al., 2018). Within each region the percentage of Il4ra expressing cells (counts greater than 0) was calculated. Additionally, for each Linnarson identified cluster within each region, the percentage of Il4ra positive cells was determined and regions of interest were visualized with pie charts.

### Accessions

Fastq files and quantified gene counts for single cell sequencing are available at the Gene Expression Omnibus (GEO) and Sequence Read Archive (SRA) under accession numbers GSE143183.

### Statistical Analysis

GraphPad Prism version 8, OriginPro 7.5, MATLAB R2017b were used for statistical analysis. All data were presented as mean ± s.e.m. N indicates the number of animals unless otherwise specified. For comparisons between two groups or more, data were analyzed by multiple *t*-test, one-way ANOVA followed by the post-hoc Tukey-Kramer test or two-way repeated measures ANOVA, as appropriate. The accepted value for significance was P <0.05. Experiments reported in this manuscript were repeated two to four times, using mice from different generations. Experimenters were blinded during data acquisition and analysis to the experimental conditions.

## Acknowledgements

We would like to thank S. Smith for editing the manuscript. We thank all the members of the Kipnis lab and the members of the Center for Brain Immunology and Glia (BIG) for their valuable comments during multiple discussions of this work. We also thank the staff of the University of Virginia Flow Cytometry Core for cell sorting and the Genome Analysis and Technology Core for library preparations and sequencing. This work was supported by grants from the National Institutes of Health (AT010416, AG034113, NS096967 and AG057496) to J.K.

## Author contributions

Z.F. Designed, performed and analyzed behavioral, brain slices preparation, dissection for snRNAseq experiments and RNAscope; T.D. and M.W. analyzed snRNAseq data; H.L. performed IHC and RNAscope experiments; A.S. performed single nucleus isolation; Z.F. and X.X. designed electrophysiology experiments; B.Z. and N.Y. performed electrophysiology experiments and analyzed the data; D.H.G. and P.H.A. assisted with IHC experiments and analysis; Z.F. analyzed RNAscope experiments; I.S. performed intra-cisterna magna injections and blinded behavior experiments; J.H. designed and performed T cell repopulation and depletion experiments, FACS, single nucleus isolation, IHC, RNAscope; provided overall advice and technical expertise; J.K. conceived the project, designed experiments, provided advice and supervised the overall work. Z.F., M.W., J.H., J.K. wrote the manuscript with input from all co-authors.

