## Supplemental Figures and legends 1-7 for "GABAergic neuronal IL-4R mediates T cell effect on memory"

### Supplementary Figure 1

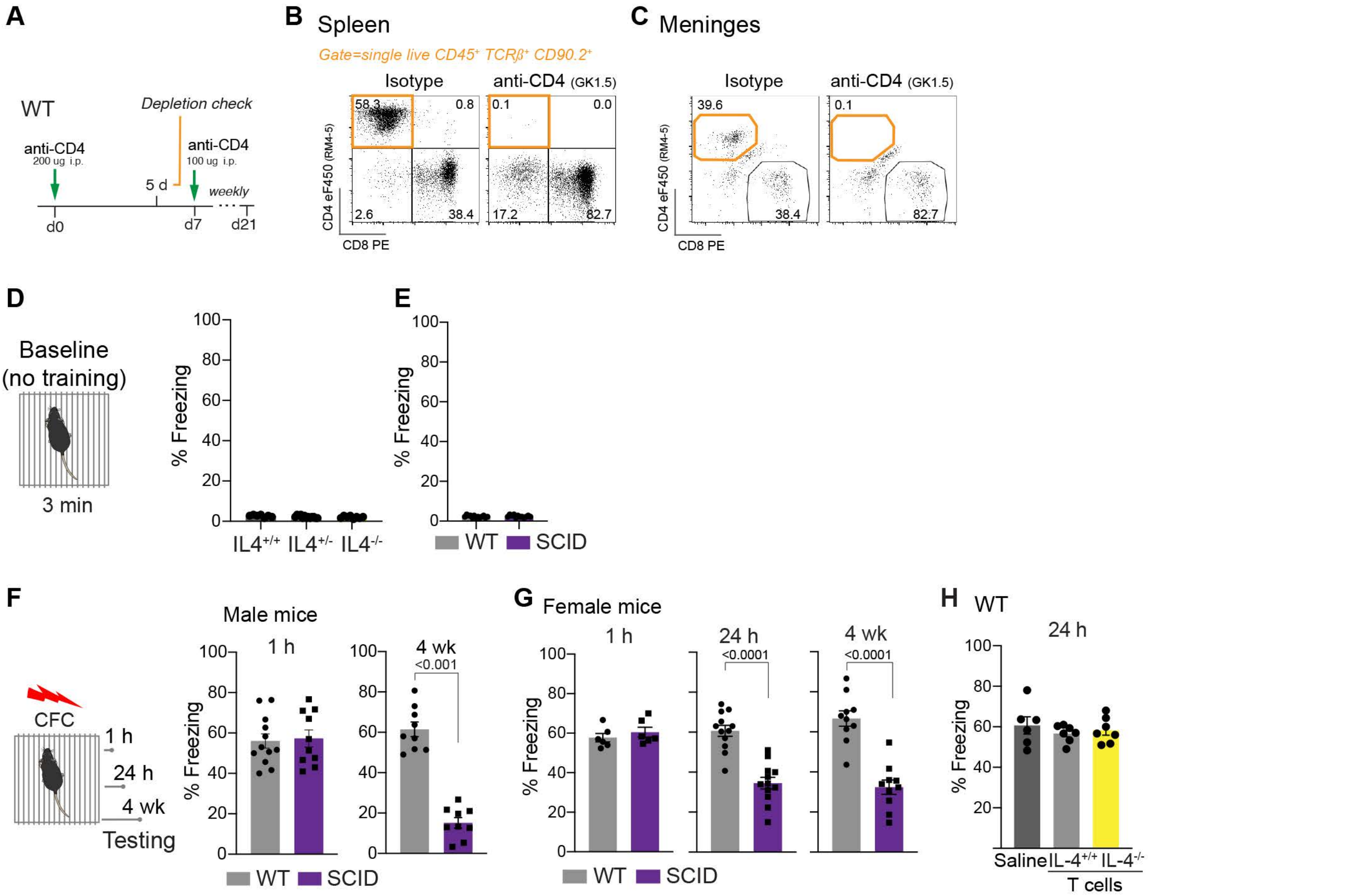

**Supplementary Data Figure 1.**

**A,** Experimental schematic for CD4 depletion of the meninges. **B**, Representative dot plot of splenic and **C**, meningeal depletion of CD4 T cells (live $CD45^+$ $TCR\beta^+$ $CD90.2^+$), 5 days after i.p. injection of 200 μg depleting antibody (clone GK1.5). N=3 mice per group. **D**, Baseline freezing level of IL-$4^{+/+}$ (n=7), IL-$4^{+/-}$ (n=13), IL-$4^{-/-}$ (n=8) mice and **E**, wildtype (WT) (n=12) and SCID (n=13) mice during habituation period in training box before fear conditioning. Each dot indicates one individual mouse. One-Way ANOVA with Tukey post hoc test. **F**, Percent freezing time of WT (n=12) and SCID (n=13) mice during 1-hour contextual fear memory testing. **G**, Adult female WT (n = 6 - 12) and SCID (n = 6 - 12) mice were trained with contextual fear conditioning and subjected to the same cage (context) for testing at different time points. Percent freezing time of wildtype 1hour, 24 hours and 4 weeks after learning. Each dot indicates one individual mouse. Two-tailed unpaired Mann-Whitney *U*-test. Data are presented as mean ± SEM. **H**, WT mice were either injected with saline or reconstituted with T cell from IL-$4^{+/+}$ mice or IL-$4^{-/-}$ mice. Percent freezing time of WT mice injected with saline (n=6), reconstituted with WT T cells (n=7) or IL-$4^{-/-}$ T cells (n=7). One-way ANOVA with Turkey post hoc test.

### Supplementary Figure 2

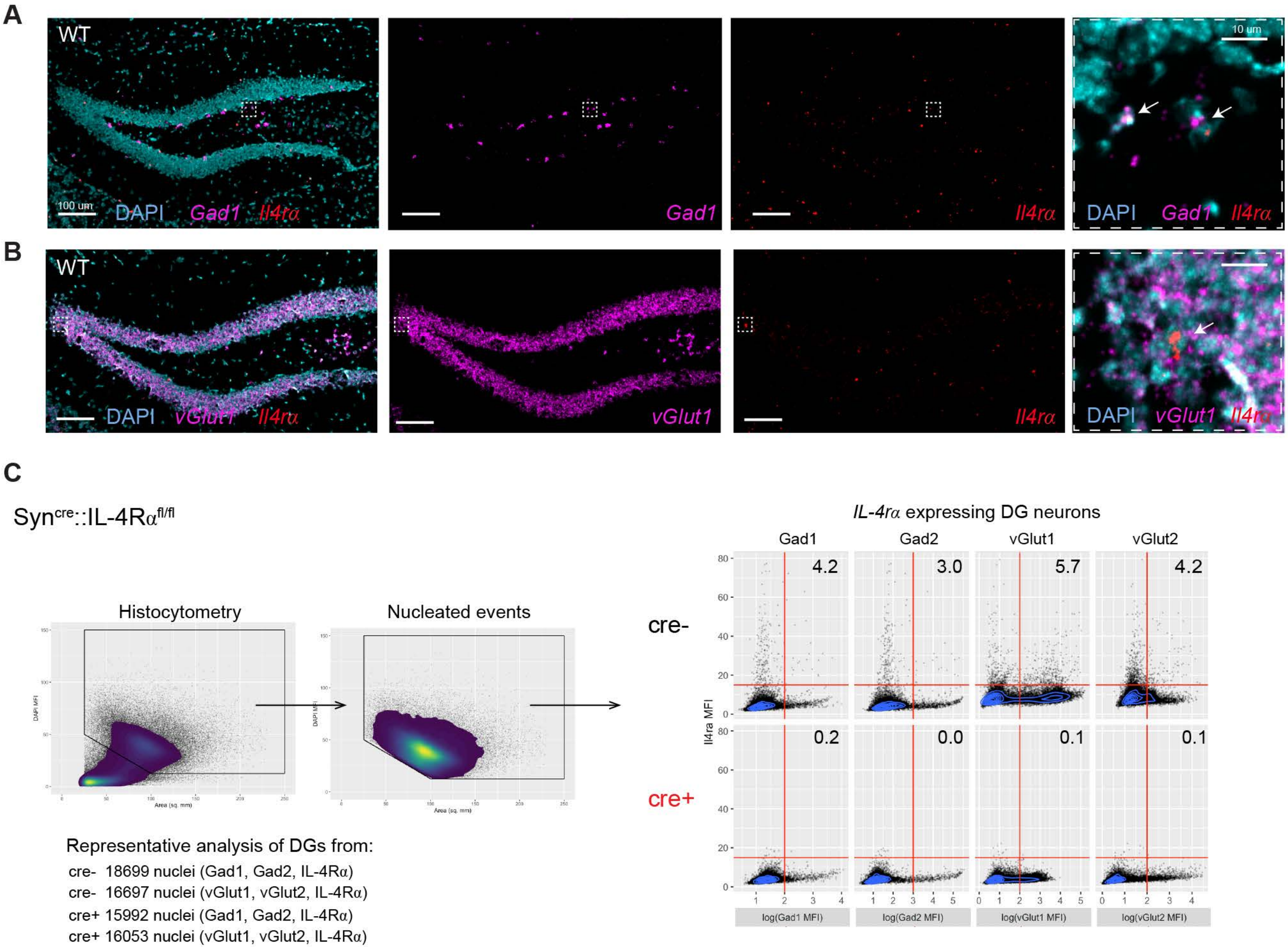

**Supplementary Data Figure 2.**

**A - B**, Representative micrographs of RNAscope showing IL-4R mRNA expression in the dentate gyrus of WT mice in (**A**) inhibitory (*Gad1*) and (**B**) excitatory (*vGlut1*) neurons. Bar = 100 µm.

**C,** Quantification of IL-4R mRNA expression in inhibitory and excitatory neurons of $Syn^{cre-}$::IL-$4Ra^{fl/fl}$ and $Syn^{cre+}$::IL-$4R^{fl/fl}$ mice. Each dot indicates one individual neuron. Representative of n= 3 mice per group.

### Supplementary Figure 3

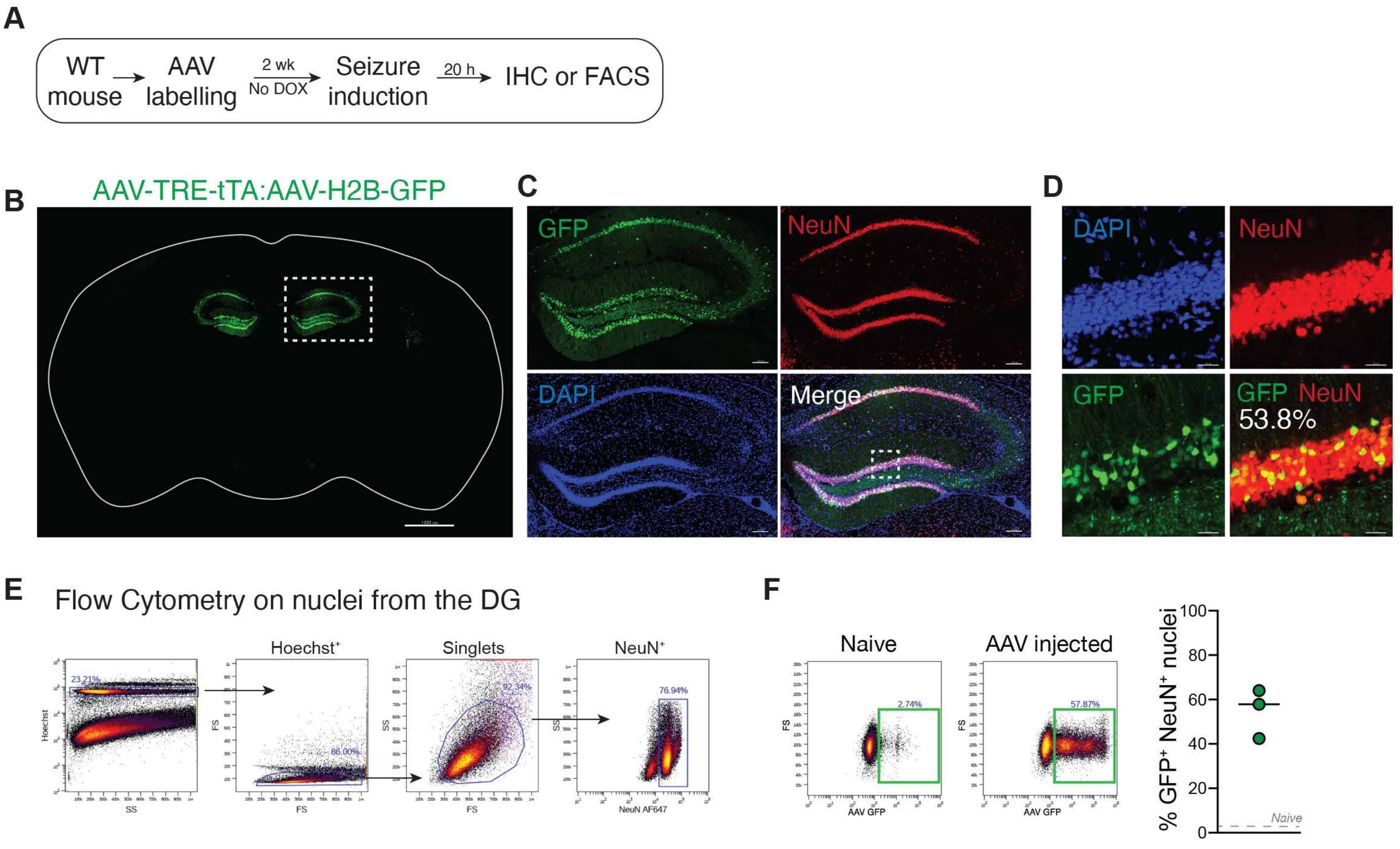

**Supplementary Data Figure 3. A,** Experimental schematic of neuronal labeling efficiency with virus-based activity labeling method. WT mice were injected with AAV-cFos-tTA and AAV-TRE-H2B-GFP virus in the hippocampus. Two weeks later mice were injected with kainic acid (25 mg per kilogram) intraperitoneally to induce seizure. Twenty hours after seizure induction, activated $GFP^+$ neurons were assessed by immunohistochemistry (B - D). **B,** Representative images of coronal brain slices 20h after seizure induction in WT mice. Bar = 1000 µm. **C,** Representative images of the hippocampus shown in **B**. Bar = 100 µm. **D,** Representative micrographs showing activated neurons in the dorsal dentate gyrus. For quantifications, the number of $GFP^+$ and $GFP^-$ $NeuN^+$ neurons was determined. Bar = 20 µm. **E,** Flow cytometry gating strategy and (**F**) statistical analysis of GFP positive neuronal nuclei after seizure induction (n=3 mice). The protocol was identical to the isolation method for snRNAseq.

### Supplementary Figure 4

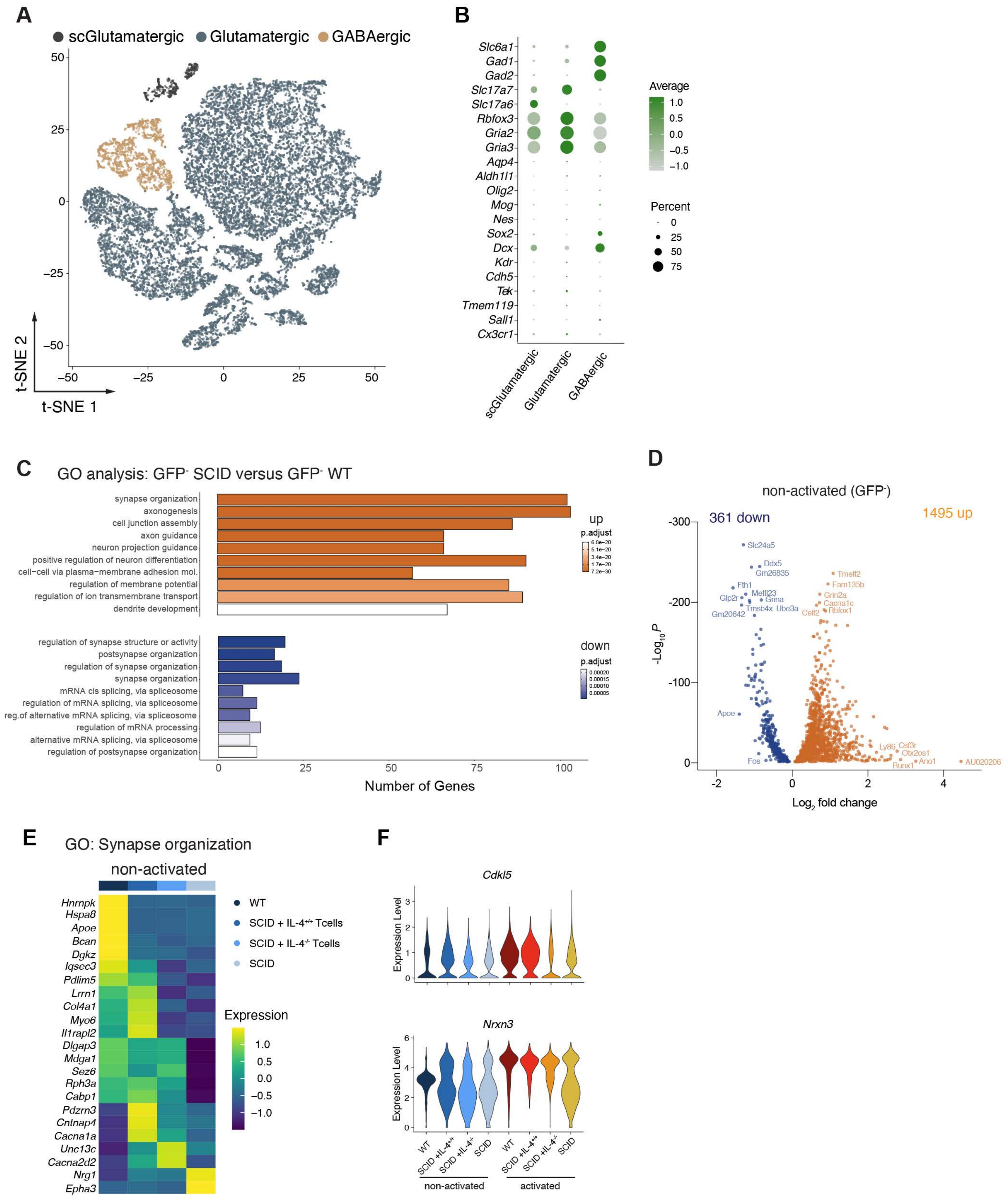

**Supplementary Data Figure 4.**

**A,** tSNE visualization of neuronal nuclei from the dentate gyrus. Colors correspond to cell type identities. **B,** Expression of markers used to identify cell cluster phenotypes. Colors corresponds to the scaled average expression of the genes within each cluster, while size corresponds to the percentage of cells from each cluster showing positive expression of the gene. **C**, Gene ontology analysis of top 10 pathways by significance comparing differentially expressed genes in non-activated $GFP^-$ neuronal nuclei from SCID and WT mice. **D**, Volcano plot showing differentially expressed genes in non-activated SCID compared to non-activated WT neurons. Top differentially expressed genes are labelled with text. **E**, Heat maps showing the average scaled expression of significantly differentially expressed genes in GO:0050808 (synapse organization) between WT, SCID with $IL4^{+/+}$ T cells, SCID with $IL\text{-}4^{-/-}$ T cells and SCID of non-activated neuronal nuclei. Mean of n=3-4 biological samples. **F**, Violin plots showing the distribution of expression of *Cdlk5* and *Nrxn3* in non-activated and activated neuronal nuclei.

### Supplementary Figure 5

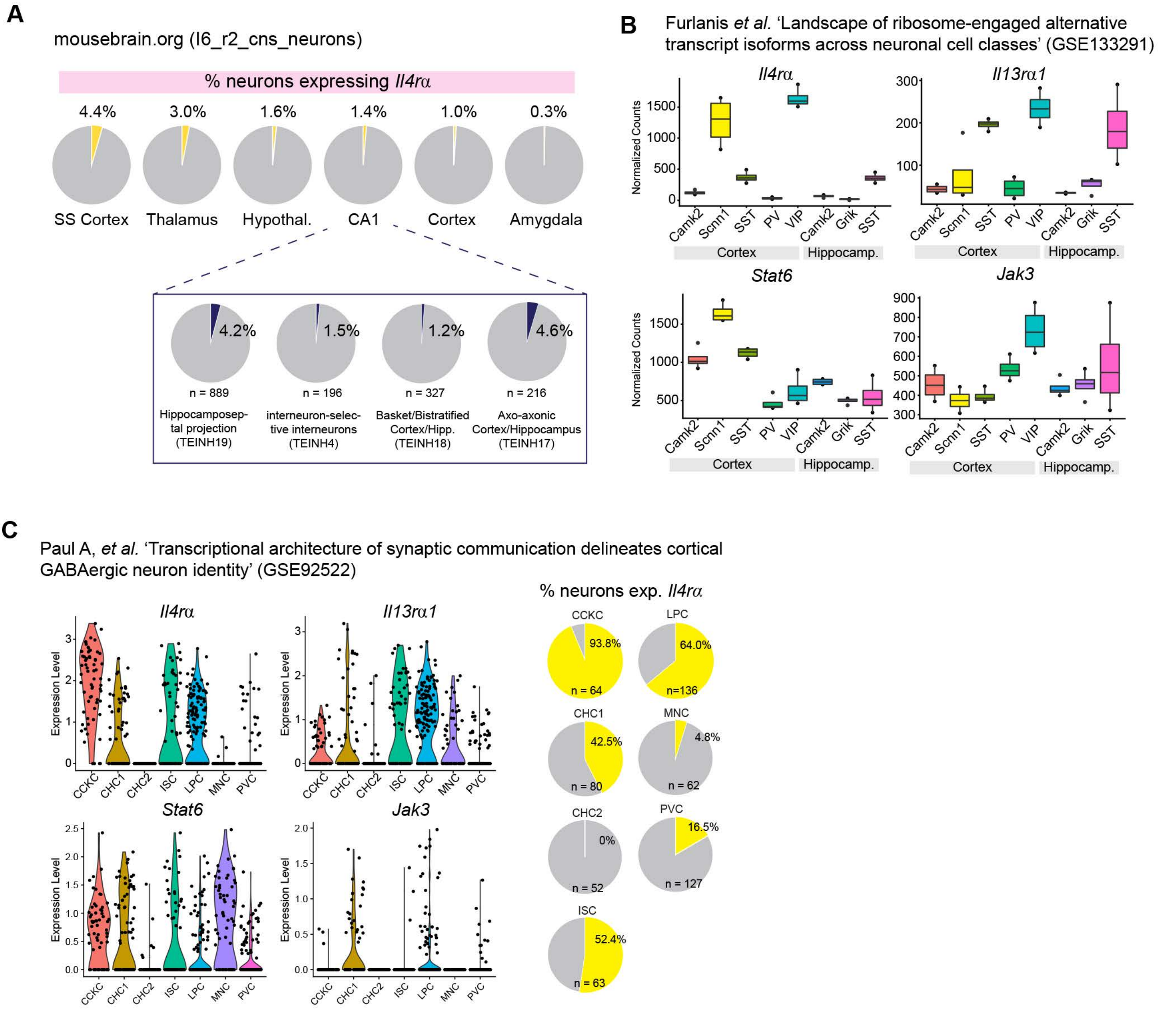

**Supplementary Data Figure 5.**

**A**, IL-4Rα mRNA expression in different brain regions from Linnarsson scRNAseq data (Amit Zeisel et al., 2018 *Cell*). **B**, Expression of *Il4rα, Il13rα1, Stat6 and Jak3* mRNA in different types of neurons in cortex and hippocampus from published RiboTAG data sets (Furlanis et al., 2019 *Nature Neuroscience*) and **C**, GABAergic neurons in cortex from scRNAseq sets (Paul et al., 2017 *Cell*).

### Supplementary Figure 6

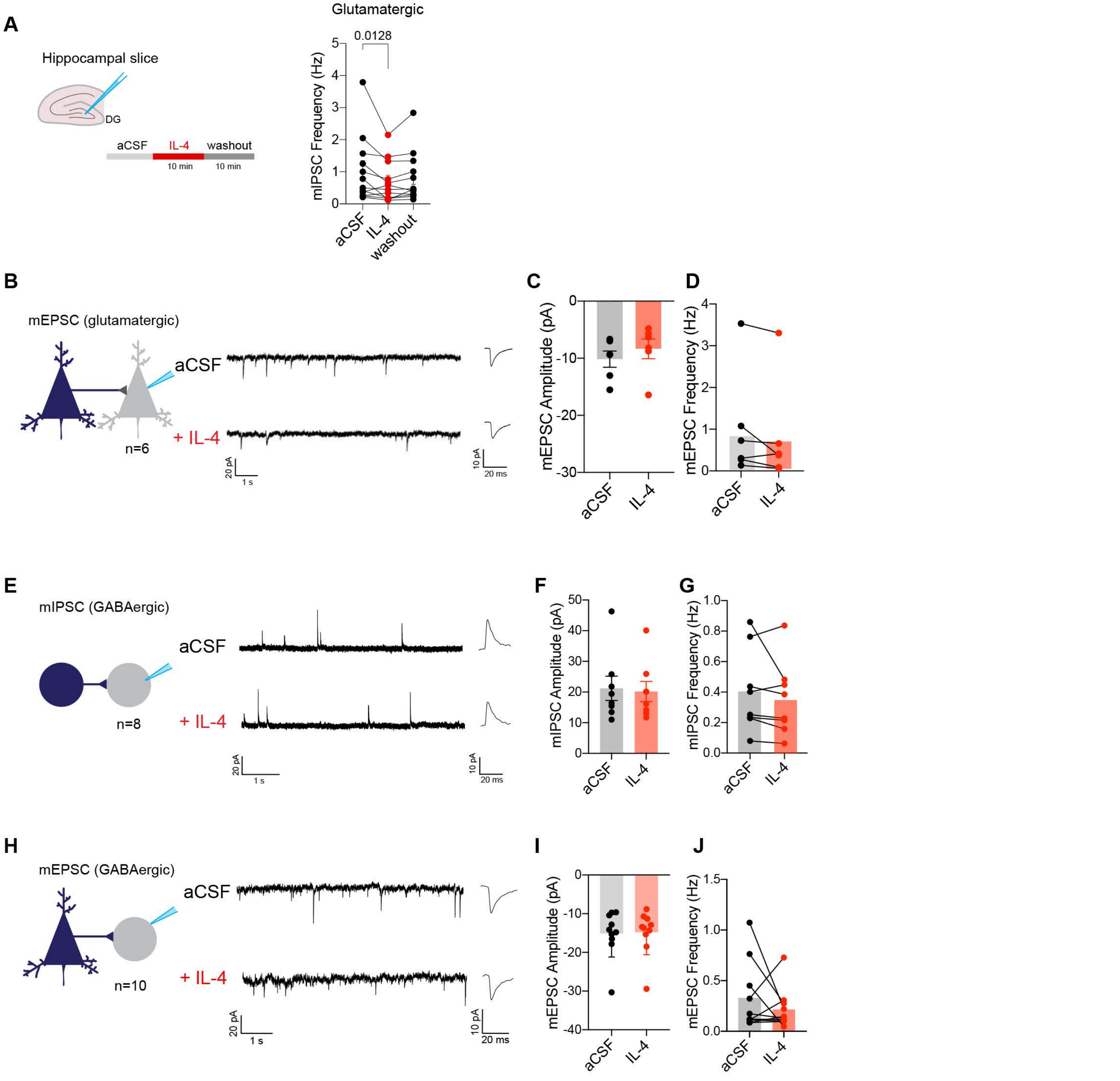

**Supplementary Data Figure 6.**

**A**, Electrophysiology recording and changes in mIPSC frequency in glutamatergic neurons before (aCSF), during and after IL-4 perfusion (washout). **B**, Example showing miniature excitatory post synaptic current (mEPSC) recording in glutamatergic neurons before and after IL-4 (100 ng/ml) perfusion. **C**, Statistical analysis of changes in mEPSC amplitude and **D,** frequency before and after IL-4 perfusion (n= 6 neurons/group). **E-G**, Example showing miniature inhibitory post synaptic current (mIPSC) recording in gabaergic neurons before and after IL-4 (100 ng/ml) perfusion (n= 8 neurons/group). **H-J**, Example showing miniature excitatory post synaptic current (mEPSC) recording in gabaergic neurons (n= 10 neurons/group). Students *t*-test. Each dot indicates one individual neuron. Data are presented as the mean ± SEM.

### Supplementary Figure 7

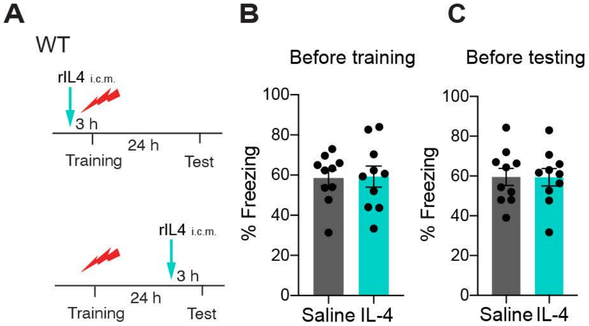

**Supplementary Data Figure 7.**

**A**, WT mice were injected with saline or 100 units of recombinant IL-4 through intra-cisterna magna (i.c.m.) 3 hours before training or testing. **B**, Percent freezing time of WT mice injected 3 hours before training. **C**, Percent freezing time of WT mice injected 3 hours before testing. Each dot indicates one individual mouse (n= 10 mice/group). Two-tailed Mann-Whitney *U*-test. Data are presented as mean ± SEM.
